# DESP: Demixing Cell State Profiles from Dynamic Bulk Molecular Measurements

**DOI:** 10.1101/2023.01.19.524460

**Authors:** Ahmed Youssef, Indranil Paul, Mark Crovella, Andrew Emili

## Abstract

There is wide interest to determine the dynamic expression of proteins and other molecules that drive phenotypic remodeling in development and pathobiology, but due to technical limitations these systems remain largely unexplored at the foundational resolution of the underlying cell states. Here, we present *DESP*, a novel algorithm that leverages independent estimates of cell state proportions, such as from single-cell RNA-sequencing or cell sorting, to resolve the relative contributions of cell states to bulk molecular measurements, most notably quantitative proteomics, recorded in parallel. We applied *DESP* to an in-vitro model of the epithelial-to-mesenchymal transition and demonstrated its ability to accurately reconstruct cell state signatures from bulk-level measurements of both the proteome and transcriptome providing insights into transient regulatory mechanisms. *DESP* provides a generalizable computational framework for modeling the relationship between bulk and single-cell molecular measurements, enabling the study of proteomes and other molecular profiles at the cell state-level using established bulk-level workflows.

## Introduction

The proteome occupies a crucial position in the functional landscape of dynamic cell states, as changes in protein expression underly biochemical processes driving phenotypic responses and altered cellular functions. In contrast to rapid advances in single-cell nucleic acid profiling, experimental challenges limit the vast majority of existing proteome studies to bulk measurements of mixed cell populations, mixing together protein abundance contributions from the individual constituent cell states and obscuring the native cellular contexts of the proteome^7,8,9^ (**Fig. 1a**). Going beyond this global birds-eye view generated by standard proteomics datasets to resolve the foundational cell-level proteomes is crucial to improve mechanistic understanding of dynamic cell states with potential to enhance the prioritization of susceptible cell subpopulations for targeted therapeutics. While the ideal scenario is to directly quantify cellular proteomes, current state-of-the-art single-cell proteomics technologies face technical challenges including the difficulty of capturing individual cells from mixtures while maintaining proteome integrity, the small amount of extracted protein, and the infeasibility of efficiently scaling up mass spectrometry runs comparable to sequencing methods^10,11^. Emerging technologies aim to address these challenges but significant limitations remain, including the extent of proteome coverage and the inability to quantify functional information such as macromolecular interactions and post-translational modifications^10,11^.

**Figure 1:**
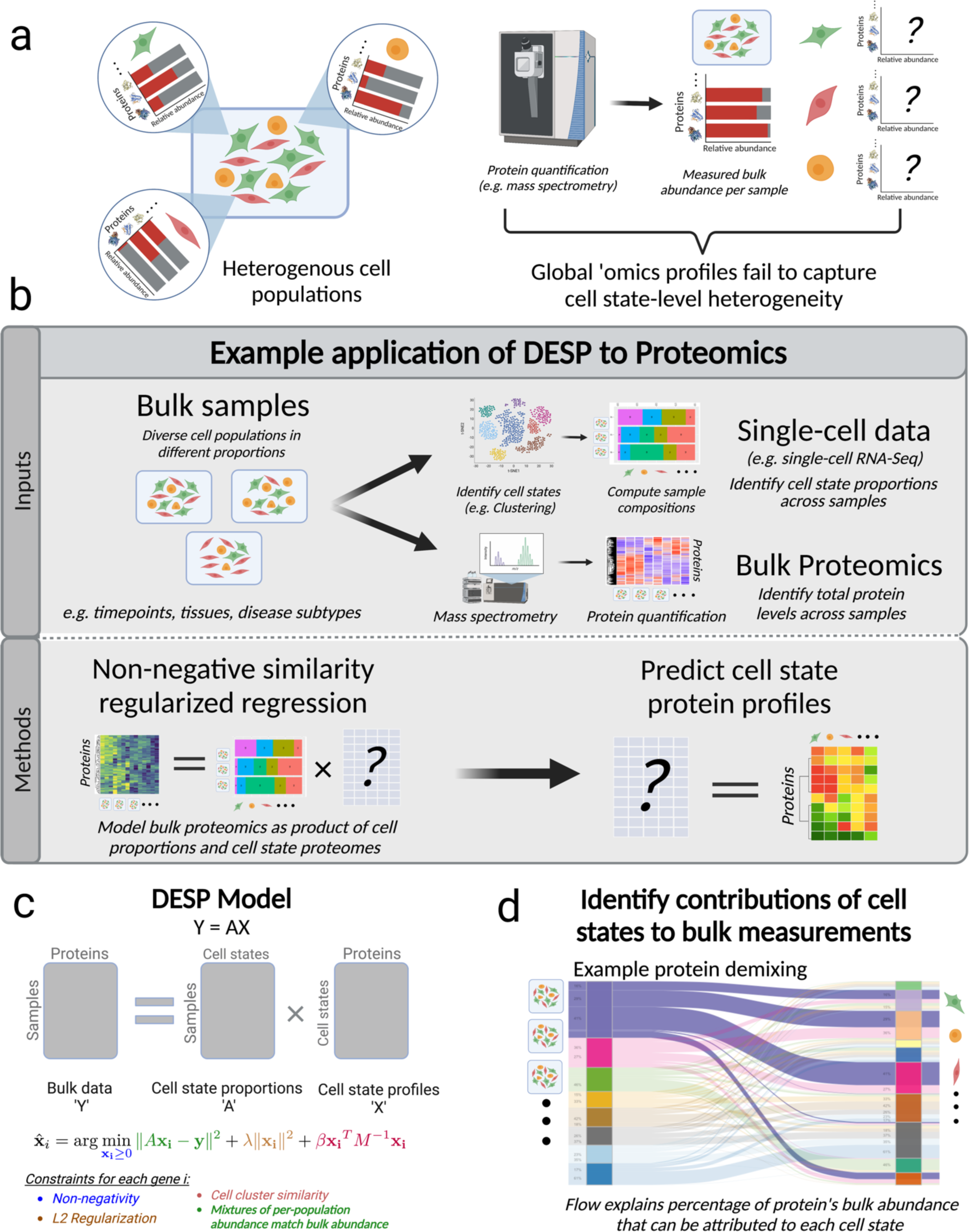
Overview of DESP algorithm. 1a: Motivation and rationale behind *DESP*. Standard ‘omic techniques such as mass spectrometry-based quantitative proteomics are often limited to bulk measurements of mixed cell populations and fail to resolve sample heterogeneity at the cell-state level. 1b: Analysis workflow. *DESP* takes as an input paired bulk omics data, e.g. proteomics, and matching single-cell data and predicts the contributions of the underlying cell states to the observed bulk measurements. 1c: Mathematical model. *DESP* models the observed bulk profiles, e.g. proteomics measurements, as the sum of unknown underlying cell state profiles. DESP applies several constraints to this formulation to predict the unknown cell state profiles. 1d: Output. Schematic of the *DESP* output showcasing its ability to demix and associate bulk sample measurements to the underlying contributing cell states for any given gene/protein.

Advances in single-cell molecular profiling, most notably single-cell RNA-sequencing (scRNA-Seq), offer a powerful approach for the identification and characterization of heterogenous cell states in dynamic biological processes, such as cell differentiation and tissue development, leading to breakthroughs in emerging domains of biomedical research^1,2^. Related computational methods can define the quantitative transcriptomic profiles underlying these discrete cell states^3^. However, these transcriptomic profiles have not proven to be accurate predictors of the corresponding protein expression patterns due to multiple biological and technical factors that lead to poor correlation between the cognate mRNA and protein levels of genes, preventing accurate inference of protein abundance using gene expression values such as those derived from scRNA-Seq^4,5,6^. This represents an important gap in our understanding of the specialized molecular programs that define cell states in heterogenous biological contexts.

As a generalizable computational solution to this challenge, we present a novel demixing algorithm titled *DESP* (<u>De</u>mixing Cell <u>S</u>tate <u>P</u>rofiles in Bulk Omics) that leverages information on cell state proportions derived from single-cell techniques to deconvolute standard bulk proteomics measurements of heterogenous cell mixtures by distinguishing the underlying contributions of the involved cell states. *DESP*’s mathematical model is designed to circumvent the poor mRNA-protein correlation, that represents a major challenge for multi-omics integration efforts, by leveraging the ability of single-cell readouts, such as scRNA-Seq or FACS, to identify the proportions of distinct cell states within dynamic heterogenous samples for which bulk proteomics measurements were generated. *DESP*’s model combines these observed changes in cell state proportions with the corresponding variations in bulk-level protein abundance to estimate the relative protein abundance levels associated with each cell state. While we demonstrate *DESP*’s utility using proteomics data in this paper, it can be applied to demix any type of bulk ‘omics data, such as metabolomics or phopshoproteomics.

We applied *DESP* to a multi-omics time-course dataset investigating the molecular changes of pre-cancerous cells undergoing TGF-Beta-induced epithelial-to-mesenchymal transition (EMT) in-vitro^12,13^. EMT is a complex biological process by which epithelial cells lose their adhesion and gradually differentiate into mobile mesenchymal cells and is of particular importance to the biomedical community due to its integral role in wound healing, tissue development, and cancer progression^12^. We analyzed bulk quantitative proteomics and scRNA-Seq data generated at eight consecutive timepoints representing MCF10A mammary epithelial cells undergoing EMT^13^, and we show that *DESP* is able to identify the corresponding cellular proteomes and recover expected patterns of canonical EMT markers. Additionally, *DESP* goes further by identifying the proteomic profile of transient cell states key to the differentiation process that were not detected at the transcriptomics level and whose signal is significantly diluted in the bulk proteomics due to the confounding presence of other cell states in the mixture. These results demonstrate a compelling use case of *DESP* while also shedding light on the transient nature of intermediate molecular mechanisms driving EMT.

## Results

### DESP algorithm overview

*DESP* is a tool for deconvoluting bulk proteomics data to the cell state-level by estimating the contributions of the underlying cell states to the observed bulk measurements. **Fig. 1b** shows an overview of *DESP* as applied to proteomics data as an example. *DESP* combines bulk proteomics data with single-cell data, such as scRNA-Seq, acquired for a set of biological samples representing different physiological contexts. Examples of potential applications for *DESP* include time-course experiments, cross-tissue ‘omics atlases, or studies comparing disease subtypes. The single-cell data is used simply to delineate the proportions of the cell states that make up the samples under study and, optionally, their quantitative similarities to each other. The resultant cell state composition matrix is fed into *DESP* alongside corresponding bulk protein measurements (**Fig. 1c**), where the objective is to find the most likely possible combination of cell state-level protein abundances whose sum would lead to the observed bulk measurements. The output of *DESP* is a matrix containing the predicted protein levels for each of the pre-defined cell states, effectively demixing the measured bulk proteomics matrix into the underlying cell state proteomes (**Fig. 1d**).

The premise of *DESP* is that the bulk proteome can be modeled as the sum of the underlying single-cell proteomes. Near-complete sequencing coverage of current single-cell technologies like scRNA-Seq make it routinely feasible to obtain the identity and proportions of cell states in given biological samples^1,2^, and advances in mass spectrometry make acquiring comprehensive measurements of bulk proteomes similarly feasible^1,2^. We thus sought to integrate these readily-available and comprehensive data types within a computational framework that could predict the experimentally-challenging single-cell proteomes. *DESP* achieves this goal by applying the *bulk-as-sum-of-single-cells* principle through modeling the measured global proteome as the product of the observed cell state proportions and the unknown corresponding cell state proteomes, providing a simple but effective formulation that can be solved mathematically to estimate the cell state proteomes (**Fig. 1c**).

The *DESP* model solves a typically under-determined problem where the number of samples is less than the number of cell states. For example, an experiment that surveys a handful of cancer subtypes will likely contain dozens of cell states involved, as such giving many possible cell-level proteomes that could lead to the observed bulk proteome. We thus apply a set of both biologically- and mathematically-motivated constraints to guide the algorithm towards the optimal solution in a deterministic manner (**Fig. 1c**). One notable constraint leverages correlations between cell state profiles in the input single-cell data to maintain the global cell state structure within *DESP*’s predictions by preserving the original state-state similarities. Notably, *DESP*’s robustness to input parameters allows it to efficiently make predictions without requiring parameter tuning (**Extended Fig. 1**). A detailed description of the mathematical model can be found in the Methods section.

*DESP* does not aim to define cell states or their proportions from ‘omics data, but rather it predicts the proteome of pre-defined cell states of interest based on the bulk proteomics data. The utility of *DESP* thus lies in the presence of established routine workflows for identifying the global proteome and cell state composition of heterogenous samples, effectively bypassing the lack of methods for deep measurement of individual cell proteomes. It is worth noting that *DESP* is agnostic of the technique used to identify cell state proportions and thus does not directly use the single-cell measurements to infer the protein levels of the given cell states, allowing *DESP* to avoid issues regarding poor correlations between different data types like RNA-Seq and proteomics.

### Multi-omics profiling of EMT

We applied *DESP* to a time-course study examining pre-cancerous cells undergoing TGF-Beta-induced EMT^13^. Bulk quantitative proteomics and scRNA-Seq data for over 5,600 genes were generated in parallel for eight consecutive timepoints as MCF10A human mammary epithelial cells underwent the EMT differentiation process (**Fig 2a**). Analysis of the scRNA-Seq data using the *Seurat* tool^14^ revealed ten discrete cell states among the 1,913 captured cells, with intermediate states emerging from the predominantly epithelial cultures before transforming into the mesenchymal end state over the timecourse (**Fig 2c,d**). The availability of both bulk protein measurements and single-cell proportion information for the samples along with the dynamic nature of the samples’ cellular compositions renders it an excellent testbed for *DESP* (**Fig. 2f**). This dataset was thus used for both algorithm validation and as a case study of a real-life application of *DESP* (**Fig 2g,h**).

**Figure 2:**
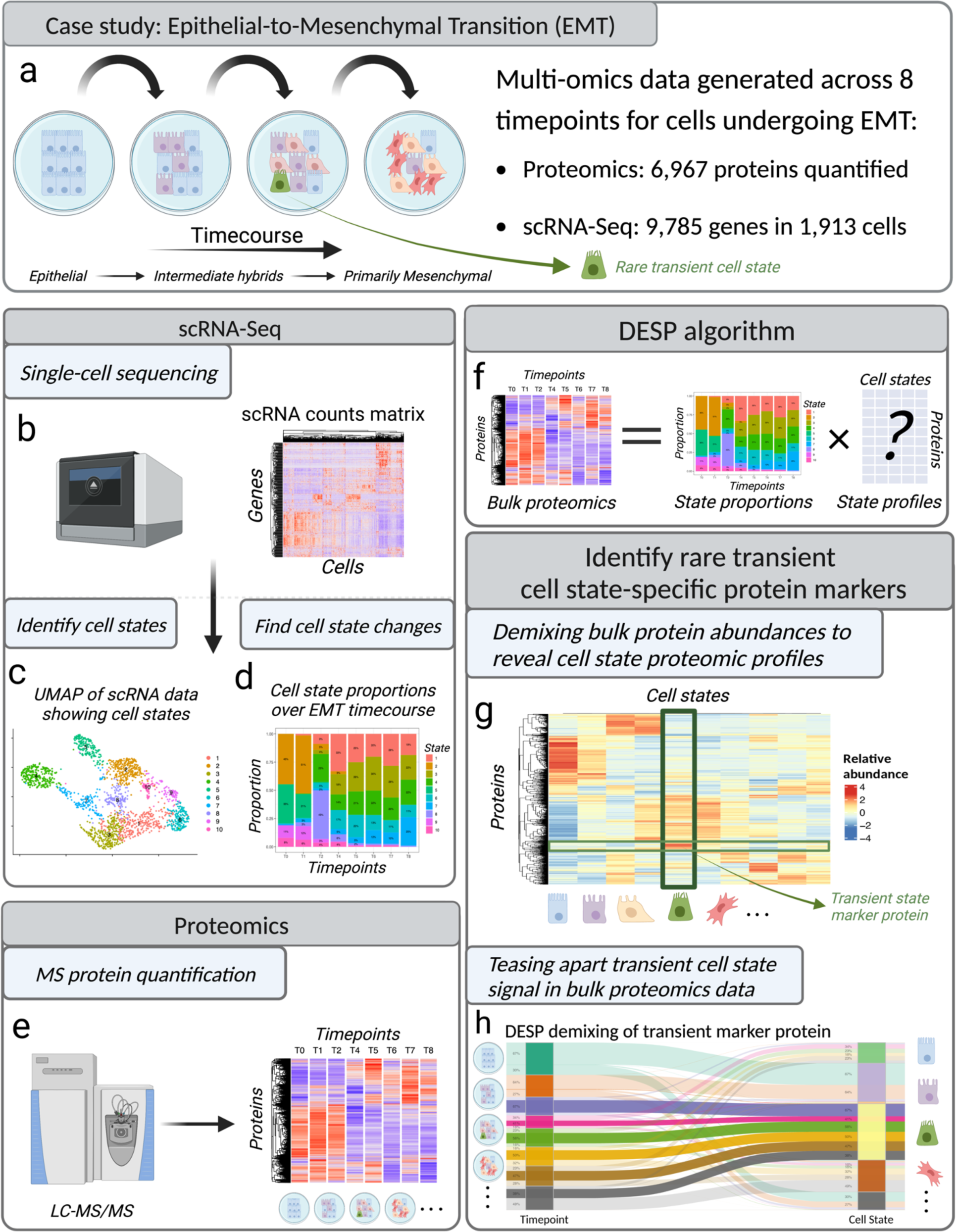
Case study: Epithelial-to-mesenchymal transition (EMT) 2a: Case study: EMT dynamic multi-omics profiling. In this experiment, bulk proteomics and scRNA-Seq data were generated for human MCF10A cells undergoing TGF-Beta-induced EMT across eight consecutive timepoints, generating molecular profiles of transient intermediate states. 2b: Dynamic scRNA-Seq profiles. Heatmap visualizes the scaled relative expression of genes (rows) across cells (columns). 2c: Identification of intermediate cell states occurring during EMT. Clustering the scRNA-Seq data reveals ten intermediate cell states as visualized on UMAP plot. 2d: Cell state fluctuations during EMT. Bar plot visualizing cell state composition at each timepoint. 2e: Mass spectrometry-based quantitative proteomics profiles. Heatmap visualizes scaled TMT reporter ion intensity, a proxy for relative abundance, of proteins (rows) across timepoints (columns). 2f: Applying *DESP* to EMT bulk proteomic profiles and scRNA-derived cell state composition matrix to predict cell state-level protein profiles. 2g: Prediction of intermediate cell state profiles using *DESP*. Heatmap visualizes the predicted relative protein abundances for each of the cell states. One putative marker of a transient cell state is highlighted as an example finding from these predictions. 2h: Identifying contribution of individual cell states to timepoint bulk proteomics measurements. Schematic shows *DESP* demixing of a given protein to distinct underlying cell states, highlighting a transient cell state signal that is lost by examining bulk measurements alone due to the presence of confounding signals from multiple cell states within each timepoint.

### Mathematical validation using pseudobulk RNA data

To validate the algorithm and assess performance, we investigated *DESP*’s ability to recover the RNA profiles of individual EMT cell states from corresponding bulk RNA profiles. We utilized the scRNA-Seq data, and not the proteomics, for the mathematical validation as it represents the ‘true’ experimentally-derived single-cell profiles to compare against *DESP*’s bulk-inferred predictions. Since *DESP*’s predictions are made at the resolution of cell states rather than single cells, we constructed representative cell state profiles from the scRNA data by averaging the expression of each gene in each cell state’s cells to represent the ground truth^27^. **Fig. 3a** shows a schematic illustrating the validation strategy, where a ‘pseudobulk’ RNA matrix is first created by summing the scRNA measurements within each of the eight timepoints. We then used *DESP* to demix these pseudobulk profiles to the level of the ten cell states present in this dataset, as guided by the scRNA-derived changes in cell state proportions across the timepoints (**Methods**). These predicted cell state profiles were then compared to the ground truth profiles.

**Figure 3:**
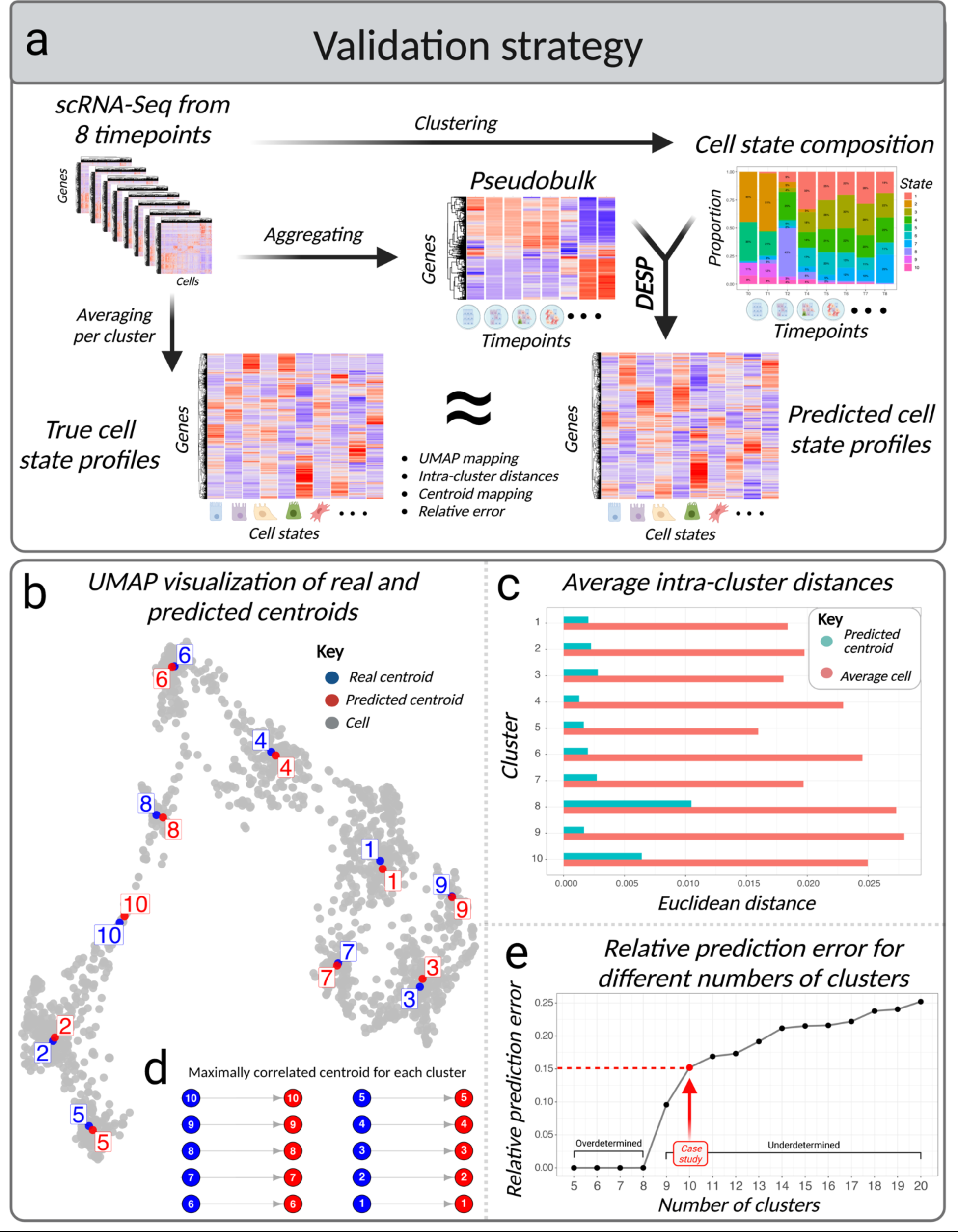
Mathematical validation of DESP using pseudobulk RNA data. 3a: Validation strategy. ‘Pseudobulk’ profiles, generated by summing the scRNA profiles in each timepoint, were demixed by *DESP* guided by the cell state compositions. The predicted *DESP* profiles were compared to the averaged per-cluster gene expression values in the input scRNA data. 3b: UMAP showing overlap of the real and predicted cell state profiles, labeled as ‘centroids’, within the global structure of the scRNA-Seq data. 3c: Bar plots showing the average Euclidean distance between each cell state’s constituent single-cell measurements to their real centroid compared to that of the predicted centroid. 3d: Mapping predicted centroids to closest real centroid based on solving the linear sum assignment problem to minimize total Euclidean distances. 3e: Relative per-gene prediction error when varying the number of cell clusters input to *DESP*, highlighting the chosen 10-cluster scenario. Standard error bars are not visible due to small error range.

Several metrics were used to compare the real and predicted cell state profiles, henceforth referred to as ‘centroids’, to determine whether *DESP*’s predictions were reasonable approximations of the ground truth. To visualize the similarity of the predicted and real centroids within the global structure of the single-cell data, we created a UMAP projection of the scRNA data with the centroids overlayed showing strong overlap between the real and predicted centroids (**Fig. 3b**). We also quantitively assessed *DESP*’s ability to recover the true centroids by computing the average Euclidean distance between each state’s constituent single-cell measurements to their real centroid compared to that of the predicted centroid, again showing that the predictions fall within the correct intra-cell state range (**Fig. 3c**). Mapping the real and predicted centroids in a one-to-one manner based on minimizing their Euclidean distances showed that *DESP* correctly assigned centroid labels with a near-zero total distance (**Fig. 3d**). Additionally, the mean per-gene relative prediction error was only 15% (IQR 6–17%), suggesting reasonable approximations of the real data even down to the level of individual genes (**Fig. 3e**).

Cell state assignments by tools such as *Seurat* can be imprecise. Hence, to ensure the robustness of *DESP* to variations in the number of input cell states, we repeated the validation with numbers of input cell states varying between 5 and 20 and found that while the prediction error expectedly increased with the number of states, it remained within a modest error range (**Fig. 3e**). We also show that *DESP* is robust to a wide range of its two input parameters with negligible effects on performance (**Extended Fig. 1**).

Since *DESP* performs predictions on a feature-by-feature basis, we also examined whether the accuracy of predictions varied with the expression of particular genes and observed that more cross-cell stable ‘housekeeping’ genes had lower prediction errors than the more variable genes, while expression magnitudes were not influential (**Extended Fig. 2a,b**). In a similar vein, we examined the relationship between the prediction errors and cell state properties and found that the larger and more homogenous states had the lowest prediction errors (**Extended Fig. 2c**).

### Mathematical validation using the Human Cell Atlas

To further validate the effectiveness of *DESP*, an additional case study was conducted using scRNA-Seq data obtained from the Human Cell Atlas^15^. The Human Cell Atlas provides a comprehensive collection of scRNA-Seq data, encompassing 48,852 genes expressed across 24 human tissues. This dataset serves as a global reference atlas of human cell expression profiles, offering an ideal opportunity to assess *DESP*’s performance on large-scale datasets (**Fig. 4a**).

For the validation process, we focused on the 36 most prevalent cell types, as determined by the number of tissues in which each cell type was present. Initially, we constructed a matrix that represented the proportion of each cell type within each tissue (**Fig. 4b**). Subsequently, we evaluated *DESP*’s ability to reconstruct the single-cell profiles by applying it to a ‘pseudobulk’ dataset. This dataset was generated by summarizing the scRNA-Seq data at the tissue level by averaging the expression of each gene within each cell type, and multiplying these averaged expressions by the corresponding cell type proportions across tissues (**Methods**).

We performed a qualitative evaluation of *DESP*’s performance by combining the real and predicted cell type profiles into one matrix and constructing a UMAP plot that visualizes these profiles in two dimensions, revealing strong overlap between the corresponding real and predicted cell type profiles within this reduced dimensional space (**Fig. 4c**). As a quantitative assessment, we computed the relative prediction error on a gene-by-gene basis by summing the absolute difference between each gene’s real and *DESP*-predicted averaged cell type profiles divided by the real cell-type profiles. The resultant median per-gene error was 15% and the mean was 20% (IQR 6 – 30%) (**Fig. 4d**). We repeated the same analysis for each cell type separately, finding a median per-cell type error of 7% and mean of 10% (IQR 2 – 13%) (**Fig. 4e**). In our experiment, it took only 6 seconds to perform the demixing for all 48,852 genes on a standard laptop machine, demonstrating *DESP*’s ability to swiftly produce accurate predictions on large-scale datasets.

### Performance comparison between transcriptomics and proteomics

*DESP* provides a mathematical framework for charting the relationship between single-cell and bulk omics datasets which depends on the assumption that the bulk data can be modeled as the sum of the single-cell data. To ensure that this relationship holds for different types of omics data, we tested the scenario where both datasets were obtained from the same samples, we applied *DESP* to matched single-cell transcriptome and protein data generated by the CITE-Seq technology from the Hao et al. study^33^. This dataset comprised 161,764 peripheral blood mononuclear cells (PBMCs) isolated from 8 individuals at three timepoints after receiving an HIV vaccine. The authors grouped the cells into 31 annotated immune cell types. Notably, there were 187 genes that had both RNA and protein measurements available for analysis.

We used *DESP* to independently demix two pseudobulk datasets created using the same underlying cell type composition matrix: the aggregated RNA data and the aggregated protein data per sample (**Methods**). Comparing the 31 *DESP*-predicted cell type profiles to the true ones measured by CITE-Seq revealed a median per-gene relative prediction error of 27% (IQR 21-34%) for the protein data, compared to 46% (IQR 28-70%) for the RNA data (**Fig. 4f**). These results indicate that *DESP* can predict single-cell protein measurements from transcriptomics-derived cell type structures with comparable, even superior, accuracy to its predictions for transcriptomics data. These observations align with previous research highlighting the consistency of high-level cell type assignments between RNA and protein measurements, despite the discordance in per-gene correlations between RNA-Seq and proteomics data^11,33^.

### Detection of transient EMT mechanisms using DESP

Applying *DESP* to demix the bulk EMT proteomics profiles allowed us to contextualize the cross-timepoint proteome changes according to the underlying cell states and characterize the distinct protein programs that occur during EMT that are lost in the bulk measurements as elaborated upon below.

To evaluate the extent to which *DESP*’s predictions provided information beyond what the bulk proteomics or scRNA data provided alone, we compared the results of performing differential expression analysis between the cell states from the *DESP* predictions versus those two original data types by themselves and found that *DESP* detected hundreds of markers for the EMT stages that were not deemed differential in the other techniques (**Extended Fig. 3**). Several of these markers are known to be involved in EMT, such as TPM2 in the E state and CD44 in the M state.

Gene set enrichment analysis also revealed more than 200 pathways that were significantly enriched among *DESP*-identified markers but not revealed by the other standard approaches, demonstrating *DESP*’s ability to glean novel insight on the involved molecular mechanisms rather than just recapitulating what existing methods would have found (**Fig. 6a**). These pathways included both canonical EMT-related pathways such as signaling, migration, and wound healing, as well as potentially novel biological processes that had not been associated with EMT before, such as cellular stress response, spliceosome assembly, and protein transport (**Fig. 6b**).

**Figure 5:**
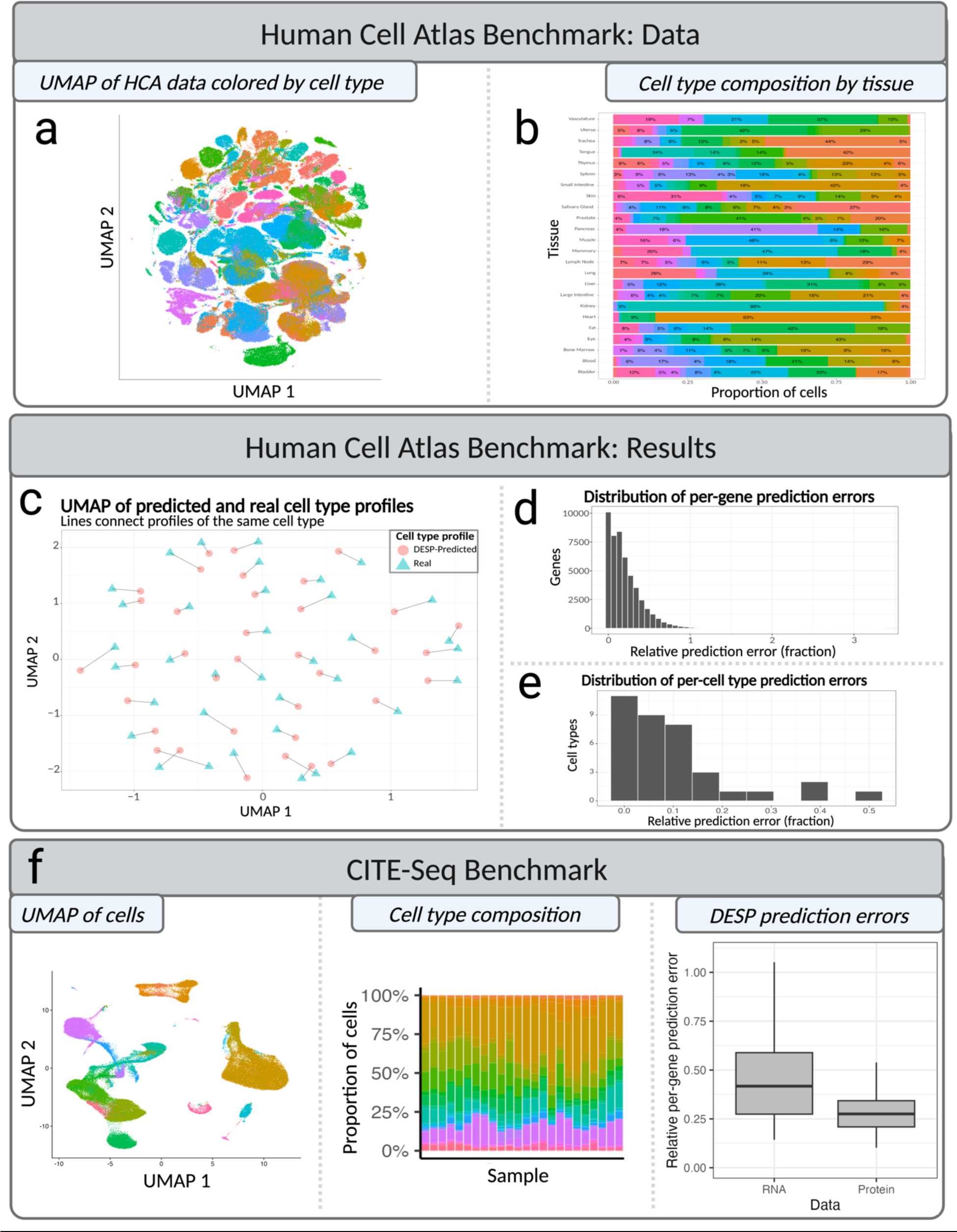
Comparing DESP predictions to experimentally-derived profiles. 5a: Dynamic EMT single-cell proteomics data used for biological validation. 5b: Heatmap of computed protein relative abundance fold-changes between cell states in single-proteomics data, scRNA data, and bulk-demixed *DESP* predictions. 5c: Pearson correlation of the protein relative abundance fold-changes between cell-states in each dataset. 5d: UMAP plot showing overlap of the real and *DESP*-predicted cell state profiles, labeled as ‘centroids’, within the global structure of the single-cell proteomics data. Inset plot: Results of mapping each predicted centroid profile to the closest real centroid profile in a one-to-one manner based on solving the linear sum assignment problem. 5e: Bar plots showing the average Euclidean distance between each cell state’s constituent single-cell measurements to their real centroid compared to that of the *DESP*-predicted centroid.

**Figure 6:**
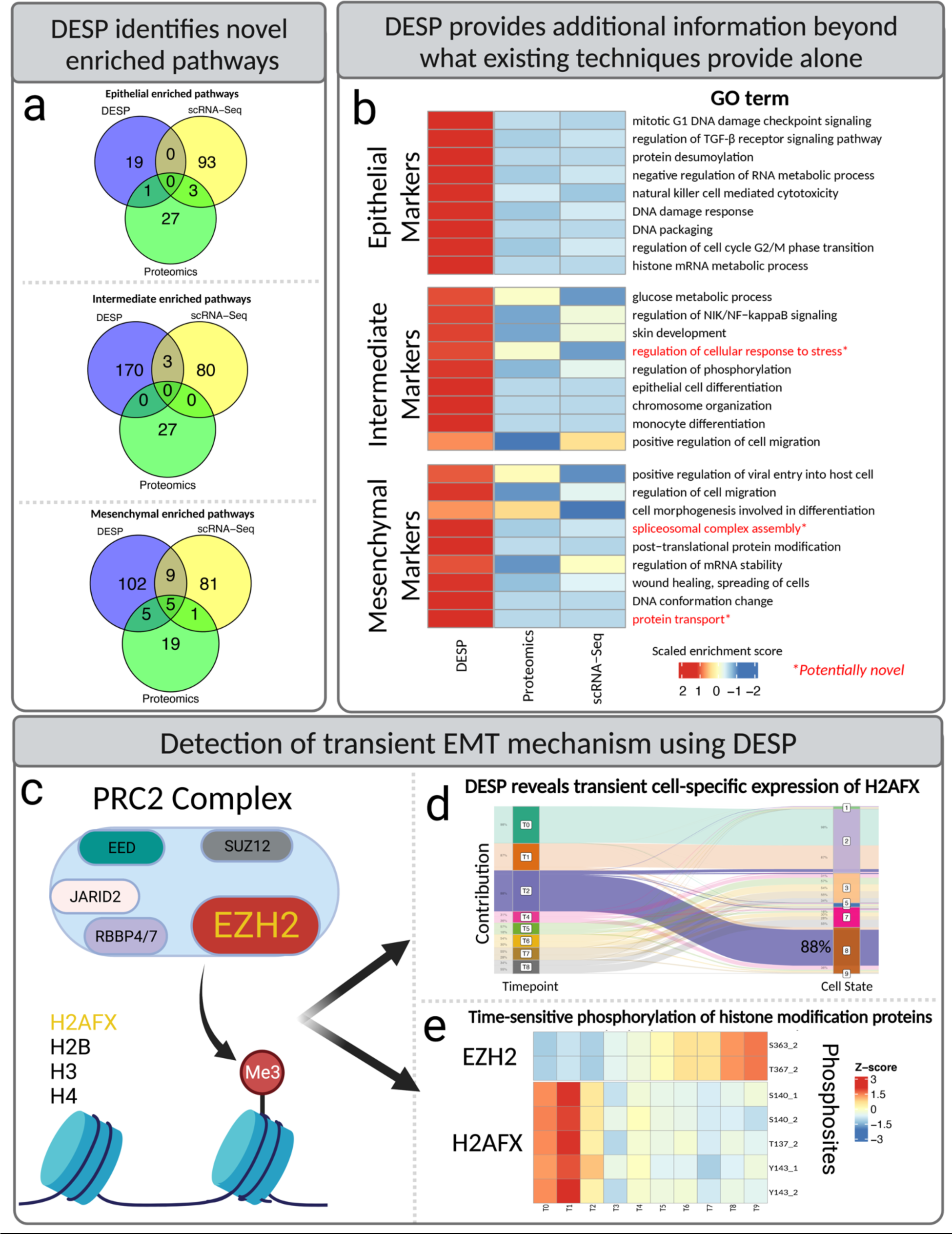
Detection of transient EMT mechanisms using DESP. 6a: Venn diagrams showing overlap of significantly-enriched pathways within each EMT stage detected by scRNA-Seq data alone, bulk proteomics data alone, and *DESP* predictions. 6b: Heatmap of relative enrichment scores of selected GO terms among cell stage markers that were enriched among *DESP*’s predictions compared to results obtained from scRNA-Seq or bulk proteomics alone. 6c: PRC2 core histone methylation complex. The EZH2 and H2AFX proteins are highlighted based on their inferred influential transient profiles detected by *DESP*. 6d: *DESP* demixing of H2AFX across transient cell states, with intermediate timepoint T2 highlighting the time-sensitive disproportionate contribution by transient cell state 8. 6e: Heatmap of temporal phosphorylation of EZH2 and H2AFX in global quantitative phosphoproteomics analysis of the same EMT samples.

The intermediate cell states are of high interest due to their transient nature and expected influential role in driving EMT. We thus sought to investigate if unique insights could be gleaned by *DESP* on the transient molecular mechanisms governing EMT. Notably, eight proteins, including three encoded by histone genes, were identified as being key to the transition due to their relative enrichment in the predicted intermediate cell state proteome compared to their diluted signal in the bulk proteomics data, representing a signal that likely would have been missed if analyzing bulk proteomics alone (**Fig. 6d**).

Of these eight proteins, H2AFX stands out as a marker (99th percentile) of the predicted intermediate proteomics profile yet with low abundance (9th percentile) in the corresponding scRNA profile suggesting extensive post-transcriptional regulation. H2AFX is a core histone protein variant that both contributes to chromatin remodeling and is post-translationally modified during EMT^17,18^. Notably, H2AFX is hyper-phosphorylated during the earliest timepoints of EMT^13^, suggesting a change in this protein’s functional activity during the initial intermediate cell states (**Fig. 6e**).

The STRING database^19^ shows that H2AFX and two other *DESP*-identified markers are functionally-linked with nine proteins annotated as EMT-associated in the GO^20^ and MSigDB Hallmark genesets^21^. One of these associated proteins is EZH2, which is part of the PRC2 chromatin remodeling complex that performs transcriptionally-inhibitory histone methylation (**Fig. 5c**). EZH2 has emerged in recent years as a promising target for cancer treatments^22,23^, with an FDA-approved drug (Tazemetostat) targeting EZH2 as a cancer treatment^24^, and another EZH2-targeting compound currently in clinical trials^25^. In contrast to its target H2AFX, EZH2 shows elevated phosphorylation during later timepoints of the experiment, suggesting the activation of a delayed regulatory ‘switch’ after three days of TGF-Beta induction.

Taken together, these findings generated by *DESP*-based proteomic demixing pinpoint specific transient events as plausible mechanisms governing EMT progression despite the RNA-protein discord.

## Discussion

Here we present, evaluate, and apply *DESP* as an algorithm for resolving the contributions of cell states to quantitative global proteomics measurements and potentially other bulk ‘omics profiles. *DESP* enables researchers to gain insights into dynamic cell state contexts from standard bulk omics workflows, helping to integrate underserved omics layers like proteomics into the rapidly evolving single-cell analysis toolkit. We validated *DESP* mathematically and obtained independent experimental evidence supporting the validity of *DESP*’s predictions. We also applied *DESP* to tease out the proteome composition of transient intermediate cell states in an in-vitro model of EMT, shining a light on key intermediate molecular mechanisms.

We note that as a new type of ‘demixing’ algorithm, *DESP* does not aim to define, nor does it require, the optimal cell state structures for the bulk data to be demixed. Rather, *DESP* solves the problem of mapping bulk omics measurements, such as proteomics, to a pre-defined cell state structure of interest to the user, independent of how that structure was originally defined. *DESP* provides a generalizable computational framework for bulk omics demixing, with planned future applications including demixing protein interaction networks and spatial transcriptomics data.

In contrast to existing omics deconvolution algorithms^26^, *DESP* does not aim to predict cell state proportions, but instead it predicts the proteome of given cell states based on demixing the bulk data. *DESP* also offers a generalizable framework applicable to all types of omics data unlike existing single-cell inference algorithms, such as TCA^35^ and BMIND^36^, that are tailored to specific data types. To facilitate adoption and usability by the community, *DESP* is available as an R package at (https://github.com/AhmedYoussef95/DESP).

## Supplementary Figures

**Extended Figure 1:**
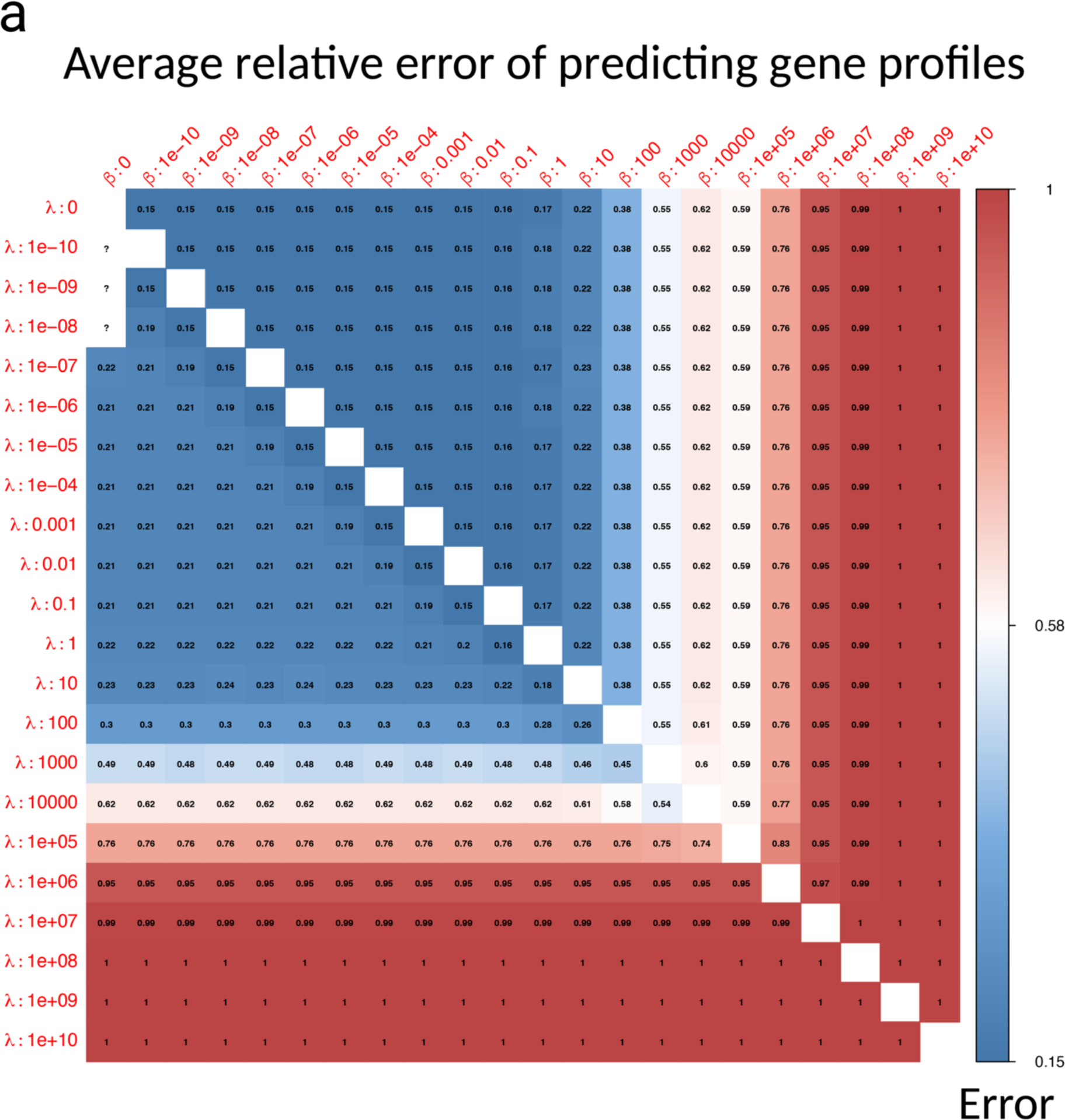
Robustness of DESP predictions to input parameters. 1a: Mean per-gene relative prediction error for applying *DESP* to EMT ‘pseudobulk’ data using a wide range of values for the two input parameters λ and β. The error is computed on a feature-by-feature basis by dividing the absolute difference between the real and predicted cell state expression values of each gene by the real expression values. Errors shown in the figure were rounded to 2 decimal points.

**Extended Figure 2:**
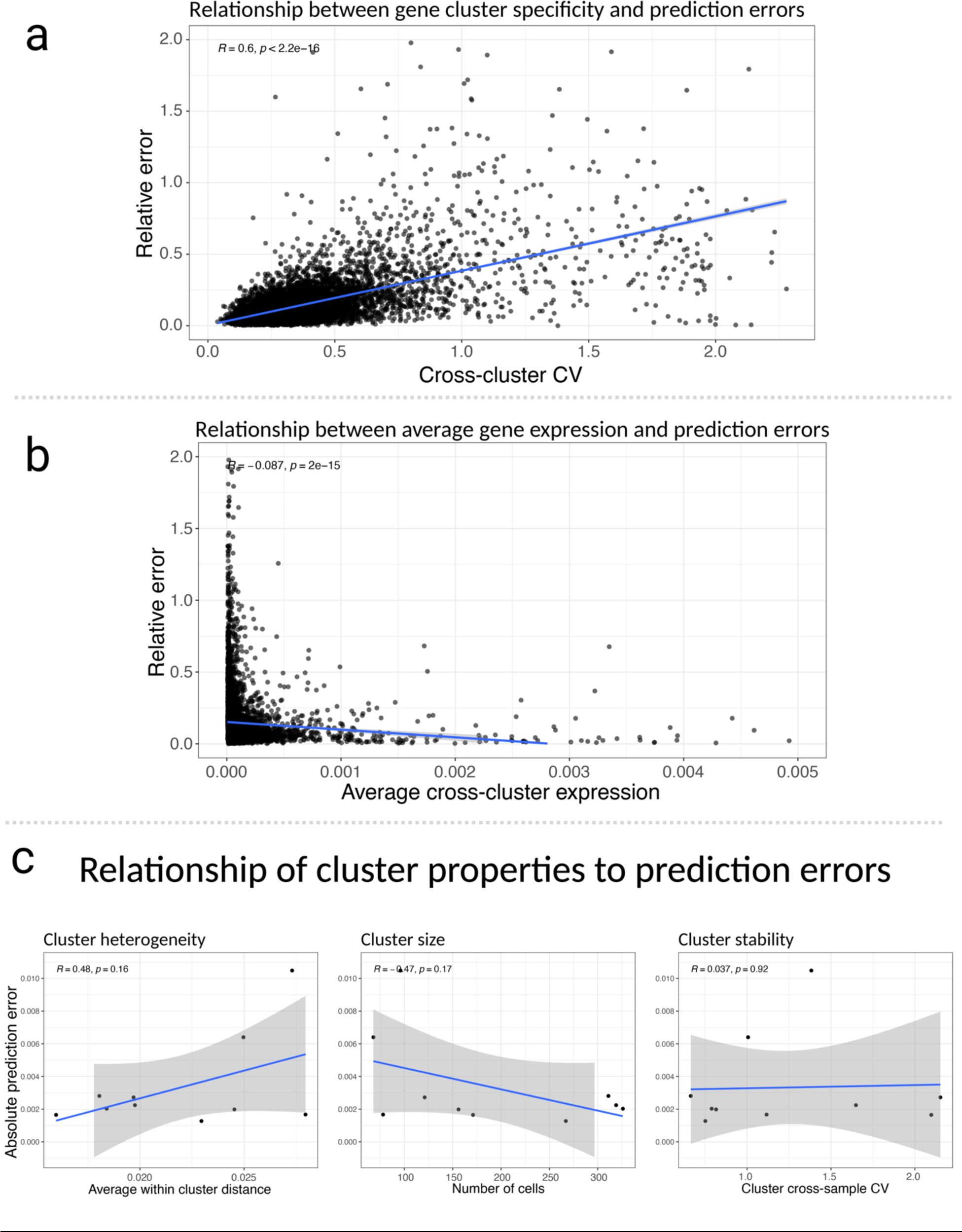
Effect of gene and cluster properties on DESP predictions. 2a: Relationship between gene cross-cluster coefficient of variation (CV) and per-gene relative prediction error. 2b: Relationship between magnitude of average gene cross-cluster expression and per-gene relative prediction error. 2c: Relationship between three mathematical properties of cell clusters (cluster heterogeneity, cluster size, and cluster proportion stability across samples) and corresponding prediction errors.

**Extended Figure 3:**
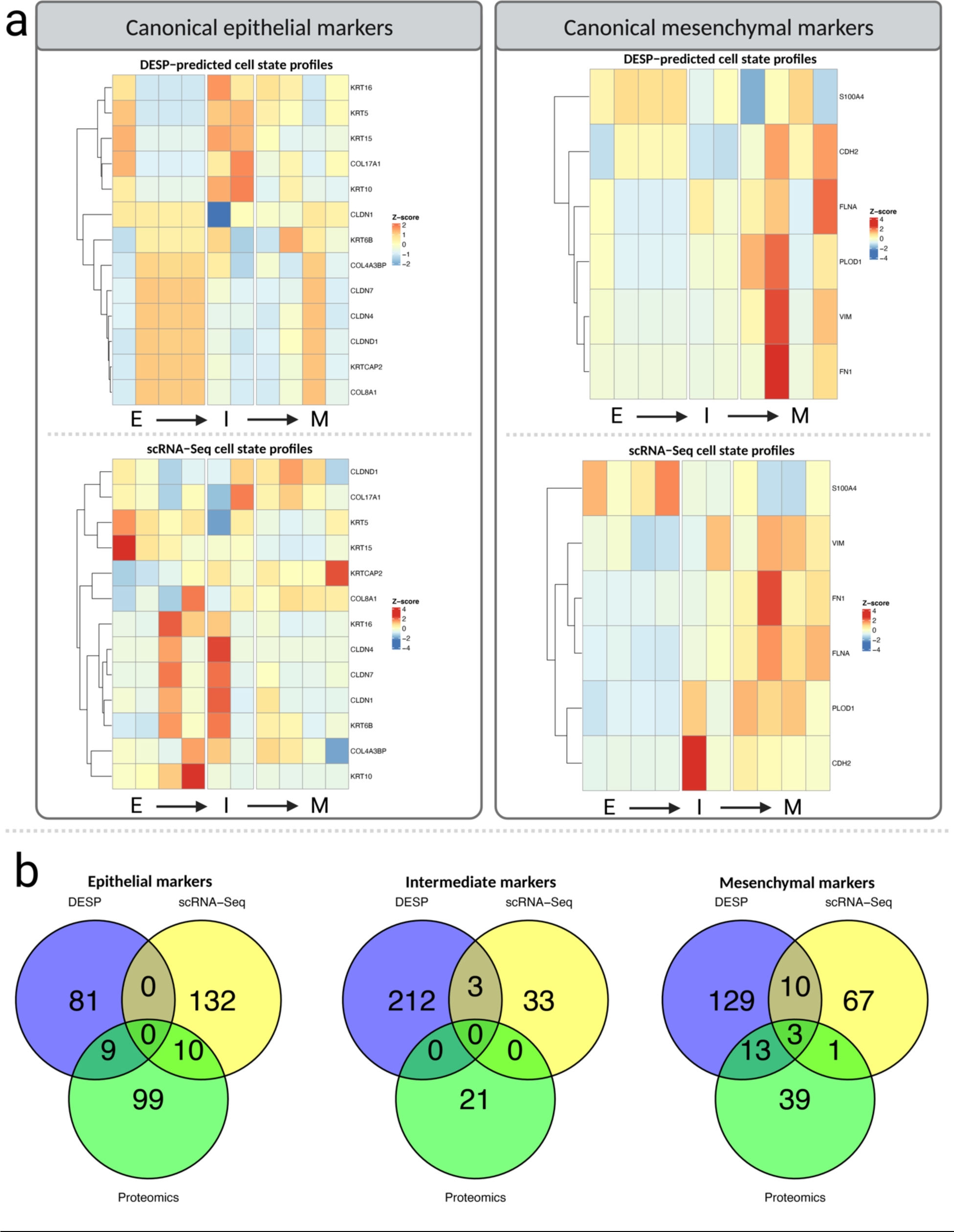
Recovery of canonical markers and discovery of novel ones. 3a: Expression/abundance patterns of selected canonical EMT markers across cell states in *DESP*’s predicted cell state protein profiles and the original scRNA-Seq data. 3b: Overlap of differentially-abundant proteins for each of the three EMT states in the scRNA-Seq data, bulk proteomics data, and *DESP*’s predicted cell state proteomics profiles.

## Methods

### DESP algorithm

#### Background

We formalize *DESP* as recovery of cell state profiles under the mixing equation *AX* = *Y*, with mixing matrix *A* (samples × states) applied to unobserved state profiles *X* (states × genes) yielding observed bulk data *Y* (samples × genes). Each pair of corresponding columns of *X* and *Y* is an independent equation; in what follows we denote a column of *X* as *x* and the corresponding column of *Y* as *y*.

This problem is challenging when there are fewer samples than states, leading to an ill-posed linear inverse problem. To constrain the solution toward meaningful and biologically likely profiles, *DESP* applies a series of regularizations as described next.

#### Nonnegativity

First, *DESP* imposes nonnegativity on the solutions. This leads to the nonnegative least squares problem:

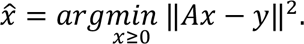

which is equivalent to the quadratic program:

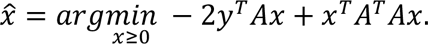

#### Tikhonov Regularization

Next, *DESP* incorporates Tikhonov regularization (ridge regression) to favor small-norm solutions:

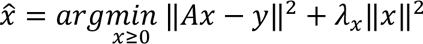

We use the notation *λ_x_* to indicate that the proper amount of regularization can be chosen based on the properties of *x*. (As shown in **Extended Figure 2**, statistical properties of individual gene expressions are correlated with estimation accuracy.) However, our results are based on using the same value of *λ_x_* for all genes. As shown in **Extended Figure 1**, using a single value of *λ_x_* across all genes yields good recovery in general with results that are relatively insensitive to the specific value of *λ_x_* (chosen within a broad range).

Expressed as a quadratic program this becomes:

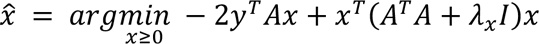

#### Similarity Regularization

Finally, we make the following observation: the process used to construct *A* may have access to additional information. For example, when scRNA clusters are used to define the cell states leading to the cluster mixtures in *A*, *DESP* can extract useful information from the cluster RNA expression profiles. We emphasize that this information can be useful even though *no assumption is made* about any particular relationship between, for example, RNA expression levels and protein abundance. *DESP* employs this strategy by incorporating a state-state similarity measure. While in principle *DESP* can use any similarity or dissimilarity matrix, in our results *DESP* operates using scRNA data as follows:

Define the matrix *C* to consist of cell state profiles with cell state profiles on the rows and genes on the columns, where entries in *C* correspond to average gene expression in the cell state. (Processing of scRNA data to form cell state profiles is discussed below.) Rows of *C* are standardized to zero mean and unit norm, to obtain *C̃*, which is used to form the correlation matrix of clusters

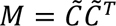

*M* is typically invertible due to the independence of cell states, so *DESP* performs similarity regularization by seeking to minimize *x^T^M*^−1^*x*

Hence to include cell state similarity as a form of regularization, *DESP* minimizes:

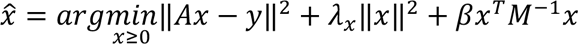

which is equivalent to the quadratic program:

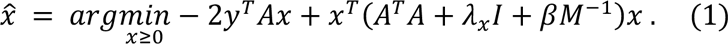

As for ***λ***_*x*_, we find that *DESP* obtains good recovery in general for ***β*** values across a fairly broad range (**Extended Fig. 1**).

#### Implementation

The R function solve.QP from the *quadprog* package can solve any of the above quadratic programs. So for each feature (e.g. gene/protein), *DESP* invokes solve.QP to solve the constrained minimization (1) to estimate cell state profiles. *DESP* also has minimal running time; in our experiments, we found that *DESP* performed predictions for >8,000 features within less than a second on a standard laptop machine.

### EMT multi-omics experiment summary

We used data from (Paul et al., 2023)^13^. In that study, cells of the human mammary epithelial cell-line MCF10A were treated with TGF-Beta to induce epithelial-to-mesenchymal transition (EMT) and samples were aliquoted at ten timepoints for multi-omics profiling. Details on the experiment and the downloadable datasets can be accessed at (Paul et al., 2023)^13^.

### Proteomics data processing

Quantitative profiles of 6,967 proteins were generated by nanoLC-MS/MS across ten timepoints (0, 4 hrs, 1–6, 8 & 12 days post-TGF-Beta injection) with three biological replicates per timepoint. Raw MS files were processed using MaxQuant (version 1.6)^32^. Tandem mass spectra were searched against the reference proteome of *Homo sapiens* (Taxonomy ID 9606) downloaded from UniProt in April 2017. Peptides of minimum seven amino acids and maximum of two missed cleavages were allowed, and a false discovery rate of 1% was used for the identification of peptides and proteins. Contaminants and reverse decoys were removed prior to downstream analysis. The average intensity across replicates was computed for each protein in each timepoint for downstream analysis. Timepoints 3 and 9 were not used in this study since they are not present in the scRNA-Seq data generated from these samples and subsequently used in downstream analysis. All data pre-processing of the MaxQuant files was performed using the R statistical software (version 4.1.0).

### Single-cell RNA-Seq data processing

Single-cell RNA-Seq quantified the expression of 9,785 genes in 1,913 cells at eight consecutive timepoints, with roughly 200 cells per timepoint. Details on the experiment and the downloadable data can be found at (Paul et al., 2023)^13^. Low-quality genes detected in less than 5% of all cells were discarded (17 genes). Lowly-expressed genes with less than three counts in at least three cells were also removed (1,240 genes). On average, each cell expressed ∼3,600 genes after filtering. Finally, the data was normalized such that each cell’s expression values sum to one. Since *DESP* ultimately combines information derived from the scRNA-Seq data with the proteomics data, we filtered the scRNA-Seq data to the set of 5,606 genes for which quantitative bulk protein measurements were also present in this dataset for the identification of co-expressed cell states prior to applying *DESP* to the proteomics data. All processing of the scRNA-Seq data was performed using the R statistical software (version 4.1.0).

### Identifying intermediate cell states from scRNA-Seq data

The cell states in the dataset were identified in an unsupervised manner based on the similarity of gene expression profiles in the scRNA-Seq data. All 1,913 cells from all the timepoints were pooled together for this analysis. The *Seurat* ^14^ R package was used to identify the cell states using their recommended workflow as follows. The data was normalized by dividing the gene counts in each cell by the total counts in the cell and multiplied by a scale factor of 10,000. The normalized counts were log-transformed after adding a pseudocount of one. The data was then centered by subtracting each gene’s expression by its average expression and scaled by dividing the centered expression levels by their standard deviations. The top 2,000 most variable genes were used to perform PCA dimensionality reduction and subsequently construct a k-nearest neighbor graph with k = 20 and 10 principal components used. Finally, the data was clustered using *Seurat*’s FindClusters function with a resolution of 1.1, defining ten co-expressed cell clusters ranging in size from 68 to 303 cells. The choice of ten clusters was made based on manual evaluation of the clustering results at different clustering resolutions. Supplementary Table 1 contains information on the markers enriched among each of these identified clusters. In the rest of the manuscript, we use the term ‘cell states’ to refer to these ten clusters.

A matrix containing the cell state profiles (denoted ‘*X*’) was constructed by averaging the expression of each gene in each cell state. A cell state proportions matrix (denoted ‘*A’)* was also constructed by counting the number of cells from each state in each timepoint. These matrices were used in the downstream validation and application of *DESP* to the EMT multi-omics data. We also tested our approach on different pre-defined numbers of states ranging between five and fifteen by varying the clustering resolution to examine the effect of varying the number of cell populations on *DESP*’s predictions (**Extended Fig 1**).

Each of the ten states identified in the previous step were also classified into one of three EMT stages based on their change in proportions across time: Epithelial (E), Mesenchymal (M), or Intermediate (I). The cell states with their maximum count in the earlier timepoints were labeled *E*, those in the later timepoints were labeled *M*, and those in the middle timepoints being *I*.

The following classifications were assigned:

- Epithelial states: 2, 5, 9
- Intermediate states: 8
- Mesenchymal states: 7

Any cell states not included in the above list were deemed not to show a distinct cross-timepoint change in proportion pattern that would best fall into one of the three biological states of interest.

### Mathematical validation using RNA pseudobulk

Prior to making inferences from the proteomics data, we first investigated *DESP*’s ability to recover the scRNA data from the bulk data at the RNA-level where we have the true single-cell profiles to compare against. To represent the bulk data that *DESP* expects as an input, we constructed a ‘pseudobulk’ matrix representing the summed gene expression levels across all cell states within each of the eight timepoints. This matrix is constructed as the product of multiplying the cell state proportions matrix ‘***A***’ and the cell state profile matrix ‘***X***’. The resultant matrix ‘***Y***’ had one value per gene in each timepoint. The cluster similarity matrix ‘***M***’ was created by computing the Pearson correlation between each pair of cell state profiles, i.e. the rows of the matrix ‘***X***’, and used to guide the algorithm towards a solution that preserves the relationship between the input cell states. We then use *DESP* to solve the under-determined problem ***Y*** = ***AX***’ to predict the cell state profile matrix ***X***’ as described in the ‘*DESP* algorithm overview’ section. Finally, the predicted cell state profile matrix ***X***’ is then compared to the real cell state profile matrix ***X*** to assess *DESP*’s performance. We utilized four different methods of assessing the prediction accuracy, each of which is detailed below.

We computed the mean per-gene relative absolute error between *X* and *X*’ by calculating the absolute difference between each gene’s real cross-state expression values in *X* and the predicted ones in *X*’ and taking the mean across all such gene-level errors. This metric was motivated by the fact that *DESP* performs its predictions independently for each gene. This metric was also used to evaluate the robustness of *DESP* to variations in its two input parameters λ and β by examining the results for different values of the parameters (Extended Fig. 1).

As a visualization of the similarity of the predicted profiles in *X*’ to the real ones in *X* within the global structure of the single-cell data, we created a UMAP projection using the R package *uwot* on the original scRNA-Seq gene expression matrix with the addition of the real and predicted cell state profiles, i.e. the rows of *X* and *X*’. The profiles were labeled on the subsequent UMAP plots to distinguish them from the individual cells (**Fig. 3b**).

To determine if *DESP*’s predictions are reasonable approximations of the real data, we also compared the predictions to the real data by computing the Euclidean distance between individual cells’ measurements and their real state average as opposed to the distance between thee predictions and the same state average. More specifically, we iterated over each of the 1,913 cells in the original single-cell expression matrix and computing the Euclidean distance between the cell’s profile and that of its cell state’s profile. Next, for each cell state we compute the average distance of its cells to the averaged cell state profile, as well as the distance between the predicted profile and the same corresponding cell state profile. We then created bar plots comparing these two distances side-by-side to determine whether the predicted profiles fall within the correct intra-state range (**Fig. 3c**).

Finally, we mapped the predicted cell state profiles to the real ones by solving the linear sum assignment problem (LSAP) using the Hungarian method as implemented in the solve_LSAP function in the *clue* ^26^ R package (**Fig. 3d**). In summary, the LSAP algorithm expects similarities between cell state profiles as the entries in the input matrix. The idea of the matching is that given two sets A and B, we find the matching that maximizes sum(similarity(a_i, b_j)) where a_i is matched with b_j, and each member of the set is matched with exactly one member of the other set. The algorithm was run twice; once with the Euclidean distance as the similarity metric for the state profiles where the algorithm minimized the sum of assigned Euclidean distances, and once with the Pearson correlation as the similarity metric where the algorithm maximized the summed correlations across cell state assignments.

### Applying DESP to demix EMT bulk proteomics data

We used *DESP* to demix the EMT bulk proteomics data down to the level of the ten scRNA-defined cell states as described in the ‘*DESP* algorithm overview’ section. The inputs to *DESP* were the processed protein-by-timepoint proteomics matrix, the cell state proportions matrix indicating the proportion of each cell state in each timepoint, and the cell state similarity matrix containing the Pearson correlations between the cell state scRNA profiles. The parameters λ and β were set to 1^−7^ and 1^−4^ respectively based on the tuning results obtained from demixing the ‘pseudobulk’ as described in the previous section. The output was a protein-by-state matrix which was used for the analysis described in the results section of the manuscript.

### Identifying cell state markers

The R package *Seurat*’s^14^ FindMarkers function was used to find the genes that distinguish each cell state in the scRNA-Seq data based on a one-vs-all differential expression analysis. An FDR cutoff of 0.05 and log fold-change cutoff of 0.5 were applied to determine the marker genes for each state. These markers are listed in **Supplementary Table 1**. This analysis was performed once at the level of the ten individual cell states, to confirm the classification of the cell states into the E/I/M stages, and once at the level of the three stages for pathway enrichment and comparing marker genes across datasets. This analysis identified 142 marker genes for the E state, 36 for the I state, and 81 for the M state in the scRNA-Seq data (**Supplementary Table 2**). For the predicted cell state proteomics profiles from *DESP*, since we have only one measurement per cell state as opposed to the multiple per-cluster cell measurements in the scRNA-Seq data, we define the marker proteins for each cell state as those with an average log2 fold-change greater than 1 between each state and the average of all the other ones (**Supplementary Table 1**). As with the scRNA-Seq data, this analysis was repeated for the three EMT stages (E, I, and M) instead of the ten cell states after the initial cell states were classified into each of the three stages. This identified 90 marker proteins for the E state, 215 for the I state, and 155 for the M state in the *DESP* predictions (**Supplementary Table 2**).

### Identifying timepoint markers in bulk proteomics

We grouped the eight timepoints in the bulk proteomics data into the E (first two timepoints), I (middle three timepoints), and M (last three timepoints) stages. Markers of each stage were defined as the proteins with an average log2 fold-change greater than 1 between each stage’s timepoints and the average of all the other ones in the bulk proteomics data (i.e. one-vs-all differential expression analysis). The data was log2-transformed and quantile-normalized prior to identifying the markers. This analysis identified 118 marker genes for the E state, 21 for the I state, and 56 for the M state in the scRNA-Seq data. These markers are listed in **Supplementary Table 2**.

### Identifying intermediate EMT markers

To focus on the mechanisms occurring exclusively during the intermediate stage, we decided to focus on the third timepoint (T2) and cell cluster 8. Cell cluster 8 is strongly-associated with this timepoint as it represents 43% of all cells in T2 but only a tiny percentage in the other timepoints. We converted the bulk protein abundances in T2 to proportions of the total T2 abundance and did the same for the predicted protein profile of cluster 8. We then divided these two percentages by each other to find the ratio between them (C8/T2), removing the few cases (∼20 proteins) where the protein has an abundance of zero in either dataset. Next, we ranked the proteins based on this ratio, and focused on proteins who have the highest/lowest ratio. The idea is that these proteins are important drivers of the transition since they are relatively important to the transient cluster 8 but whose signal seems diluted in the bulk proteomics based on this ratio, probably due to the presence of other clusters contributing to the bulk signal. The resultant ratios had a mean and median of ∼1, and we decided to focus on the 99th percentile, representing the 70 proteins whose ratio was above 1.32. In parallel, we also took the 208 proteins defined as upregulated markers of the predicted protein profile of cell cluster 8 based on a one-vs-all differential expression analysis as a second set of intermediate markers. The two analyses described above thus produced two sets of proteins of interest for their inferred role in the transitional phase of EMT, and we found eight proteins that overlapped between them. The results section of the manuscript elaborates on some of the inferred roles of these proteins in the context of EMT.

### Recovering associations with EMT-related genes

We curated a set of 320 EMT-associated genes and interrogated the set of predicted intermediate markers for physical and/or functional associations with those genes. The EMT-associated genes came from two sources: 200 genes from the MSigDB Hallmark geneset^21^, and 132 genes associated with at least one of the 11 GO Biological Process terms with “epithelial mesenchymal transition” in the title^20^. 12 genes overlapped between the two sets, while none of our predicted markers were present in this set, suggesting potential novelty. We used the STRING database^19^ to check for reported physical interactions and/or functional associations between our marker genes and the EMT-associated genes. To cast a wide net, we considered associations reported based on all of STRING’s scoring methods, such as RNA co-expression and experimentally-determined interactions, but filtered them to include only high-confidence interactions (STRING combined score > 70%).

### Pathway enrichment analysis

We used the *enrichR*^29^ tool to identify biological pathways significantly enriched for each cell state’s protein and RNA markers as well as the bulk proteomics timepoint markers based on a hypergeometric test with a p-value cutoff of 0.01. The pathway databases interrogated included GO Biological Process, GO Molecular Function, and GO Cellular Component from the 2021 version of the GO database^20^, in addition to MSigDB^21^, OMIM^30^, and KEGG^31^. We also used the *enrichR* tool to compute and compare the enrichment scores of gene sets of interest across the markers (**Fig. 5b, Extended Fig. 5**). *enrichR*’s combined score, computed as log(p) * z where p is the Fisher test p-value and z is the z-score for deviation from expected rank, was used for this analysis as a measure of the relative enrichment of a given geneset among a given set of cell state markers. **Supplementary Table 3** lists the detected significantly-enriched pathways from the *DESP* predictions, scRNA-Seq, and bulk proteomics data.

### Phosphoproteomics data processing

Phosphoproteomics data for the same set of cells was downloaded from (Paul et al.)^13^, with three replicates per timepoint. The dataset consists of phosphorylation data for 8,727 phosphosites across 1,776 unique proteins. The intensity was averaged across replicates for each phosphosite.

### Human Cell Atlas case study

Publicly-available scRNA-Seq data was downloaded from the Human Cell Atlas^15^. The data consisted of gene expression values for 58,870 genes across 24 tissues encompassing 177 annotated cell types. The following pre-processing steps were applied to the *raw* scRNA-Seq counts:

1. Removed genes with zero variance across cells (removed 1,619 genes)
2. Removed lowly-expressed genes by retaining genes with more than 3 counts in at least 3 cells (removed 8,399 genes)
3. Normalized the data such that each cell sums to 1
4. Retained the top 36 cell types based on the number of tissues each cell type is expressed in (led to a minimum of 3 out of 24 tissues per cell type)

To perform *DESP* demixing, we constructed a matrix with the average expression of each gene in each cell type. This matrix represented the true cell type profiles, i.e. the ground truth to compare *DESP*’s predictions against. We also constructed a matrix containing the proportion of each cell type among the cells belonging to each tissue (**Fig. 4b**). We then applied *DESP* to these two matrices, using a λ parameter value of 0 and β value of 1^−10^, to predict the expression of the 48,852 retained genes across the 36 cell types.

### Analysis of CITE-Seq data

Publicly-available CITE-Seq data was downloaded from Hao et al^33^. The dataset consisted of parallel RNA and protein measurements for 161,764 PBMCs isolated from 8 volunteers at three timepoints (days 0, 3, and 7) after receiving an HIV vaccine, with a set of 187 overlapping genes between the RNA and protein measurements. Gene mapping between the RNA and protein measurements were carried out based on common Ensembl identifiers^34^. The authors assigned cell types at 3 hierarchical levels, and we decided to focus our analysis on the intermediate level which grouped the cells into 31 annotated immune cell types. The single-cell data was normalized using the *Seurat* package such that each cell sums to 10,000. We then summarized both the RNA and protein data into the following matrices:

- *A* (cell type composition): Number of cells from each cell type in each sample. This is consistent for both the RNA and protein data.
- *X* (cell type profiles): average expression of each gene/protein in each cell type.
- *Y* (pseudobulk): Matrix multiplication of *A* and *X*, representing aggregated single-cell data in each sample.

We quantitatively evaluated *DESP*’s ability to predict the cell type profiles in the matrix *X* given the *A* and *Y* matrices, comparing performance for predicting the RNA and protein profiles separately. Since *DESP* predictions are performed on a feature-by-feature basis, we computed the prediction error as the absolute relative difference between the true and predicted cell state values for each feature, and created boxplots comparing *DESP*’s per-gene relative prediction error for the RNA and protein data separately (**Fig. 4f**).

### Supplementary Text

#### Testing robustness of DESP predictions with different numbers of cell states

In the EMT case study detailed in the manuscript, we identified ten transitional cell states occurring during EMT. These cell states were identified by clustering the single-cell RNA-Seq data to reveal groups of transcriptionally-correlated cells using the *Seurat* R package (**Methods**). Since determining the optimal number of cell states form scRNA-Seq data is not an objective process, we repeated our mathematical validation of *DESP* using different numbers of input cell states to examine the robustness of *DESP*’s predictions.

To vary the number of cell states, we used different clustering resolutions within *Seurat*’s FindClusters algorithm followed by repeating the process for constructing a cell state proportions matrix detailing the proportion of each cell state within each of the timepoints. We tested *DESP* using numbers of clusters varying between 5 and 20 and computed mean per-gene relative prediction error for each case (**Fig 3e**). For cases where there were 8 or less cell states, the problem was overdetermined due to the presence of eight timepoints in the data and thus the prediction error was zero. Starting from nine clusters and above, the problem is underdetermined meaning that multiple possible combinations of cell state protein profiles could lead to the observed pseudobulk measurements. As expected, we found that the prediction error increased as the number of cell states increased, with the error ranging from 10-25% for cases with 9-20 clusters respectively.

#### Testing robustness of DESP predictions to input parameters

*DESP* expects two input parameters denoted as λ and β (**Methods**). In the EMT case study described in the manuscript, we set the values of these parameters to 1^−7^ and 1^−4^ respectively. To test the robustness of *DESP*’s predictions to variations in the values of the input parameters, we repeated the mathematical validation of *DESP* using combinations of the input parameters ranging from 1^−10^ to 1^10^, with increasing steps of an order of magnitude each, for each of the two parameters and quantified the mean per-gene relative prediction error for deconvoluting the scRNA-Seq ‘pseudobulk’ data (**Extended Fig. 1**).

We found that the error remained stable at 15% for a wide range of combinations of the input parameters, with predictions made with values of β ranging between 1^−7^ and 0.01 and corresponding values of λ of the same or less value all giving nearly identical results with errors of 15%. Furthermore, the error never exceeded 22% for any combination of the parameters with values between 1^−10^ and 1 for each parameter. The prediction errors were smaller for deconvoluting the pseudobulk single-cell proteomics data (**Fig. 5d,e**), with prediction errors at 2% for any value of β between 1^−10^ and 0.1 and a corresponding λ of the same or less value, and never exceeding 16% for any combination of values between 1^−10^ and 1 for either parameter. These results demonstrate the robustness of *DESP* to variations in the input parameters and that there is no need to hypertune the input parameters for different input datasets.

#### Effect of gene/protein properties and cell cluster properties on DESP’s predictions

*DESP* performs predictions on a feature-by-feature basis. Depending on the input data, these features will typically be genes or proteins. We examined the correlation between mathematical properties of the features in the bulk data and their corresponding prediction error based on applying *DESP* to the scRNA-derived pseudobulk matrix as described in the mathematical validation chapter of the Methods section. To explore the relationship between properties of the genes and their corresponding predicted values, for each of the 8,528 genes in the dataset we computed the following measures:

- ***Expression specificity:*** This metric looked at the relative specificity of a given gene’s expression to the clusters. The coefficient of variation (CV) of the gene’s cluster-specific expression values is computed as the standard deviation of the per-cluster expression values divided by their mean. Genes with a lower CV can be interpreted as being relatively stable ‘housekeeping’ genes due to the smaller variance in their expressions across clusters.
- ***Average expression:*** This metric is concerned with the relative quantity of each gene’s transcripts and is simply computed as the mean expression of the gene across clusters.

Each of the above properties was correlated to the gene’s relative prediction error, defined as the absolute difference between the real and predicted per-cluster values divided by the real value. A Pearson correlation coefficient of 0.6 was observed between the per-gene prediction errors and their expression specificity, indicating that the less variable ‘housekeeping’ genes tended to have higher prediction accuracy (**Extended Fig. 2a**). Meanwhile, the Pearson correlation coefficient was only −0.01 with the genes’ average expression values, suggesting that the magnitude of a gene’s expression is less influential to *DESP*’s predictions (**Extended Fig. 2b**).

In a similar vein, we computed the Pearson correlation coefficient between the following mathematical properties of the cell clusters and their corresponding prediction errors measured as the Euclidean distance between the predicted centroids and the real ones:

- ***Cluster heterogeneity***: For each cluster, the average Euclidean distance between each member cell’s scRNA expression profile and that of its corresponding cluster centroid, i.e. the average expression profile of all cells belonging to the cluster.
- ***Cluster size***: The number of cells within each cluster.
- ***Cluster fluctuation across samples***: The coefficient of variation of each cluster’s proportion in each timepoint.

We found a positive correlation between cluster heterogeneity and the prediction error (Pearson R^2^ = 0.48) indicating that the prediction error was lower for the more homogenous clusters, a negative correlation with size (Pearson R^2^ = −0.47) suggesting better prediction for larger clusters, and near-zero correlation with cross-sample cluster fluctuations (Pearson R^2^ = 0.04) (Extended Fig. 2c, d, e). These results are potentially confounded however by the fact that cluster size and heterogeneity were strongly negatively-correlated to each other (Pearson R^2^ = −0.65).

## Acknowledgements

AE and MC acknowledge a generous pilot award from the Rafik B. Hariri Institute for Computing and Computational Science & Engineering at Boston University. We acknowledge the generous data and advice from Dr. Nikolai Slavov, Dr. Anand Asthagiri, and Saad Khan at Northeastern University. We acknowledge the advice and helpful discussions with Dr. Josh Campbell, Dr. Stefano Monti, and Dr. Trevor Siggers at Boston University.

## Author Contributions

AY, AE, and MC conceived the project. AY and MC performed the computational analysis. IP provided biological expertise. AY wrote the manuscript. All authors revised the manuscript. AE and MC supervised the project.

## References

1. Ogbeide, Silvia, et al. “Into the Multiverse: Advances in Single-Cell Multiomic Profiling.” Trends in Genetics, vol. 38, no. 8, 2022, pp. 831–843., 10.1016/j.tig.2022.03.015.

2. Stuart, Tim, and Rahul Satija. “Integrative Single-Cell Analysis.” Nature Reviews Genetics, vol. 20, no. 5, 2019, pp. 257–272., 10.1038/s41576-019-0093-7.

3. Chen, Geng, et al. “Single-Cell RNA-Seq Technologies and Related Computational Data Analysis.” Frontiers in Genetics, vol. 10, 2019, 10.3389/fgene.2019.00317.

4. Buccitelli, Christopher, and Matthias Selbach. “MRNAs, Proteins and the Emerging Principles of Gene Expression Control.” Nature Reviews Genetics, vol. 21, no. 10, 2020, pp. 630–644., 10.1038/s41576-020-0258-4.

5. Fortelny, Nikolaus, et al. "Can we predict protein from mRNA levels?." Nature 547.7664 (2017): E19.

6. Brion, Christian, et al. “Simultaneous Quantification of Mrna and Protein in Single Cells Reveals Post-Transcriptional Effects of Genetic Variation.” ELife, vol. 9, 2020, 10.7554/elife.60645.

7. Aebersold, Ruedi, and Matthias Mann. “Mass-Spectrometric Exploration of Proteome Structure and Function.” Nature, vol. 537, no. 7620, 2016, pp. 347–355., 10.1038/nature19949.

8. Uhlen, M., et al. “Tissue-Based Map of the Human Proteome.” Science, vol. 347, no. 6220, 22 Jan. 2015, pp. 1260419–1260419, 10.1126/science.1260419.

9. Marx, Vivien. “A Dream of Single-Cell Proteomics.” Nature Methods, vol. 16, no. 9, 2019, pp. 809–812., 10.1038/s41592-019-0540-6.

10. Specht, Harrison, et al. “Single-Cell Proteomic and Transcriptomic Analysis of Macrophage Heterogeneity Using scope2.” Genome Biology, vol. 22, no. 1, 2021, 10.1186/s13059-021-02267-5.

11. Brunner, Andreas-David, et al. “Ultra-High Sensitivity Mass Spectrometry Quantifies Single-Cell Proteome Changes upon Perturbation.” Molecular Systems Biology, vol. 18, no. 3, 2022, 10.15252/msb.202110798.

12. Lamouille, Samy, et al. “Molecular Mechanisms of Epithelial–Mesenchymal Transition.” Nature Reviews Molecular Cell Biology, vol. 15, no. 3, 2014, pp. 178–196., 10.1038/nrm3758.

13. Paul, Indranil, et al. “Parallelized multidimensional analytic framework applied to mammary epithelial cells uncovers regulatory principles in EMT.” Nature Communications, 14(1), 688, 2023, 10.1038/s41467-023-36122-x.

14. Satija, Rahul, et al. “Spatial Reconstruction of Single-Cell Gene Expression Data.” Nature Biotechnology, vol. 33, no. 5, 2015, pp. 495–502., 10.1038/nbt.3192

15. Regev, Aviv, et al. “The Human Cell Atlas.” ELife, vol. 6, 2017, 10.7554/elife.27041.

16. Slavov, Asthagiri, et al. (in preparation)

17. Segelle, Alexandre, et al. “Histone Marks Regulate the Epithelial-to-Mesenchymal Transition via Alternative Splicing.” Cell Reports, vol. 38, no. 7, 2022, p. 110357., 10.1016/j.celrep.2022.110357.

18. Lone, Imtiaz Nisar, et al. “The Role of Histone Variants in the Epithelial-to-Mesenchymal Transition.” Cells, vol. 9, no. 11, 2020, p. 2499., 10.3390/cells9112499.

19. Szklarczyk, Damian, et al. “The String Database in 2021: Customizable Protein–Protein Networks, and Functional Characterization of User-Uploaded Gene/Measurement Sets.” Nucleic Acids Research, vol. 49, no. D1, 2020, 10.1093/nar/gkaa1074.

20. Gene Ontology Consortium. The Gene Ontology resource: enriching a GOld mine. Nucleic Acids Res. 2021 Jan 8;49(D1):D325–D334. doi: 10.1093/nar/gkaa1113. PMID: 33290552; PMCID: PMC7779012.

21. Liberzon, Arthur, et al. “The Molecular Signatures Database Hallmark Gene Set Collection.” Cell Systems, vol. 1, no. 6, 2015, pp. 417–425., 10.1016/j.cels.2015.12.004.

22. Kim, Kimberly H, and Charles W Roberts. “Targeting EZH2 in Cancer.” Nature Medicine, vol. 22, no. 2, 2016, pp. 128–134., 10.1038/nm.4036.

23. Duan, Ran, et al. “EZH2: A Novel Target for Cancer Treatment.” Journal of Hematology & Oncology, vol. 13, no. 1, 2020, 10.1186/s13045-020-00937-8.

24. Straining, PharmD, Rachael, and William Eighmy, PharmD. “Tazemetostat: EZH2 Inhibitor.” Journal of the Advanced Practitioner in Oncology, vol. 13, no. 2, 2022, pp. 158–163., 10.6004/jadpro.2022.13.2.7.

25. Wishart DS, Feunang YD, Guo AC, Lo EJ, Marcu A, Grant JR, Sajed T, Johnson D, Li C, Sayeeda Z, Assempour N, Iynkkaran I, Liu Y, Maciejewski A, Gale N, Wilson A, Chin L, Cummings R, Le D, Pon A, Knox C, Wilson M. DrugBank 5.0: a major update to the DrugBank database for 2018. Nucleic Acids Res. 2017 Nov 8. doi: 10.1093/nar/gkx1037.

26. Avila Cobos, Francisco, et al. “Benchmarking of Cell Type Deconvolution Pipelines for Transcriptomics Data.” Nature Communications, vol. 11, no. 1, 2020, 10.1038/s41467-020-19015-1.

27. Bilous, Mariia, et al. “Metacells Untangle Large and Complex Single-Cell Transcriptome Networks.” BMC Bioinformatics, vol. 23, no. 1, 2022, 10.1186/s12859-022-04861-1.

28. Hornik, Kurt. “A Clue for Cluster Ensembles.” Journal of Statistical Software, vol. 14, no. 12, 2005, 10.18637/jss.v014.i12.

29. Kuleshov, Maxim V., et al. “ENRICHR: A Comprehensive Gene Set Enrichment Analysis Web Server 2016 Update.” Nucleic Acids Research, vol. 44, no. W1, 2016, 10.1093/nar/gkw377.

30. Amberger, Joanna S., et al. “OMIM.org: Online Mendelian Inheritance in Man (OMIM®), an Online Catalog of Human Genes and Genetic Disorders.” Nucleic Acids Research, vol. 43, no. D1, 2014, 10.1093/nar/gku1205.

31. Kanehisa, Minoru, et al. “Kegg as a Reference Resource for Gene and Protein Annotation.” Nucleic Acids Research, vol. 44, no. D1, 2015, 10.1093/nar/gkv1070.

32. Tyanova, Stefka, et al. “The Maxquant Computational Platform for Mass Spectrometry-Based Shotgun Proteomics.” Nature Protocols, vol. 11, no. 12, 2016, pp. 2301–2319., 10.1038/nprot.2016.136.

33. Hao, Yuhan et al. “Integrated analysis of multimodal single-cell data.” Cell vol. 184,13 (2021): 3573–3587.e29. doi:10.1016/j.cell.2021.04.048

34. Fiona Cunningham, et al., Ensembl 2022, Nucleic Acids Research, Volume 50, Issue D1, 7 January 2022, Pages D988–D995, 10.1093/nar/gkab1049

35. Rahmani, E., Schweiger, R., Rhead, B. et al. Cell-type-specific resolution epigenetics without the need for cell sorting or single-cell biology. Nature Communications 10, 3417 (2019). 10.1038/s41467-019-11052-9

36. Wang, Jiebiao, Kathryn Roeder, and Bernie Devlin. "Bayesian estimation of cell type-specific gene expression with prior derived from single-cell data." Genome Research (2021) 31: 1807–1818.

